# Precise base editing for the *in vivo* study of developmental signaling and human pathologies in zebrafish

**DOI:** 10.1101/2020.12.12.422520

**Authors:** Marion Rosello, Juliette Vougny, François Czarny, Maria Caterina Mione, Jean-Paul Concordet, Shahad Albadri, Filippo Del Bene

## Abstract

While zebrafish is emerging as a new model system to study human diseases, an efficient methodology to generate precise point mutations at high efficiency is still lacking. Here we show that base editors can generate C-to-T point mutations with high efficiencies without other unwanted on-target mutations. In addition, we established a new editor variant recognizing an NAA PAM, expanding the base editing possibilities in zebrafish. Using these approaches, we first generated a base change in the ctnnb1 gene, mimicking oncogenic mutations of the human gene known to result in constitutive activation of endogenous Wnt signaling. Additionally, we precisely targeted several cancer-associated genes among which cbl. With this last target we created a new zebrafish dwarfism model. Together our findings expand the potential of zebrafish as a model system allowing new approaches for the endogenous modulation of cell signaling pathways and the generation of precise models of human genetic disease associated-mutations.

## INTRODUCTION

With the recent technological advances in precise gene editing, the use of zebrafish in genetic engineering studies has drastically increased in the last five years (Patton and Tobin, 2019; Santoriello and Zon, 2012). The CRISPR (Clustered Regularly Interspaced Short Palindromic Repeats)/Cas9 system is indeed a remarkably powerful gene editing tool (Sander and Joung, 2014) that enables the rapid and efficient generation of loss-of-function mutations in this model. This system relies on the specific binding of a sgRNA-Cas9 complex that initially interacts with DNA through the NGG Protospacer Adjacent Motif (PAM) sequence and is next stabilized by annealing of the sgRNA to the 20 base pairs (bp) upstream of the PAM which triggers the Cas9 protein to introduce a double-strand break (DSB). This technique is nowadays widely used in zebrafish notably to produce knock-out alleles (Hwang et al., 2013b). However, although successful in some studies (Armstrong et al., 2016; Bai et al., 2020; Hisano et al., 2015; Hruscha et al., 2013; Hwang et al., 2013a; Irion et al., 2014; Zhang et al., 2016), precise modifications and knock-in by homology-directed repair (HDR) remain inefficient in this animal model (Albadri et al., 2017).

Recently, a CRISPR/Cas9-based technology has been developed to precisely edit single bases of DNA without introducing DSBs in human cells (Koblan et al., 2018; Komor et al., 2016; Komor et al., 2017). The method is based on the fusion of a Cas9-D10A nickase with a cytidine deaminase giving rise to a Cytidine Base Editor (CBE). CBE converts C to T bases in a restricted window of 13 to 19 nucleotides (nt) upstream of the PAM sequence (Fig. 1A). In zebrafish, a CBE has been shown to work but efficiencies were limited, with less than 29% editing being induced and, in most cases, at least 5% of unwanted INDEL (Insertion or Deletion) mutations were detected (Carrington et al., 2020; Zhang et al., 2017). For these reasons, these editing strategies have not been so far favored by the zebrafish community and no direct CBE applications have been performed to the study of developmental processes or to generate human disease models. However, since this first generation of CBEs, several studies in cell culture have optimized and engineered new base editor variants with increased gene editing efficiency reaching up to 90% and the elimination of undesired INDEL mutations (Koblan et al., 2018). Recent progress has also been made on the generation of CBEs able to recognize other PAM sequences, allowing to broaden CBE editing possibilities (Jakimo et al., 2018; Koblan et al., 2018).

**Fig. 1.**
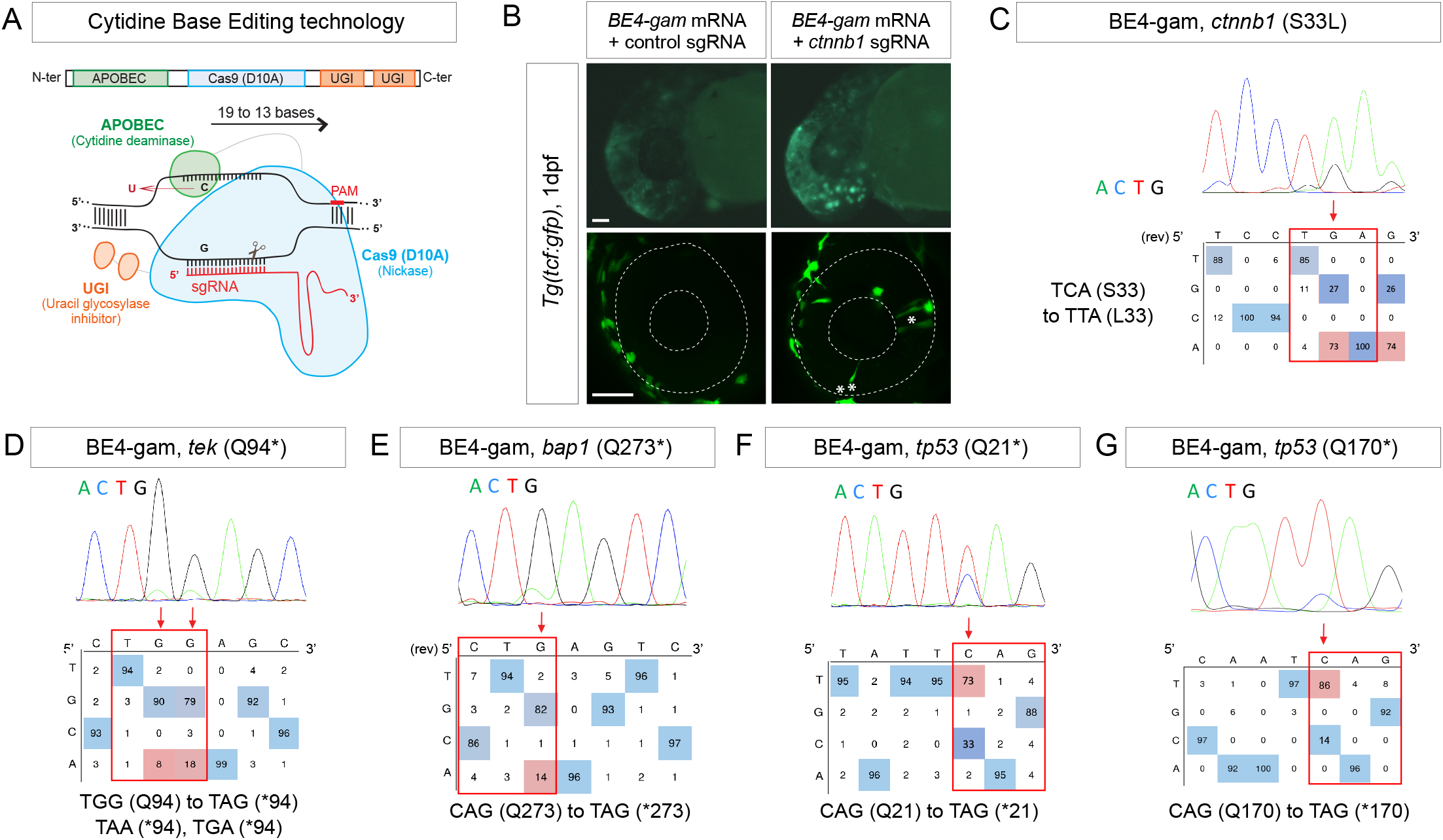
Efficient endogenous activation of Wnt signaling pathway and tumor suppressor genes targeting using BE4-gam in zebrafish. (A) Schematic representation of the Cytidine Base Editor technology. (B) Activation of Wnt signaling via S33L mutation in β-catenin. 1dpf *Tg(tcf:gfp)* embryo injected with *BE4-gam* mRNA and *ctnnb1 (S33L)* sgRNA or control scrambled sequence. The upper panel shows an overall increase of GFP -positive cells upon the injection of the *BE4-gam* mRNA and *ctnnb1 (S33L)* sgRNA compared to the control situation. The lower panel shows maximal z-projection of lateral view of the injected embryos where ectopic GFP signal in retinal progenitor cells (white stars) can be detected whereas control embryos do not show any fluorescence in the retina at this stage. (C-G) DNA sequencing chromatogram of targeted loci with the BE4-gam and obtained C-to-T conversion efficiencies. (C) S33L mutation in β-catenin upon C-to-T conversion in *ctnnb1* reached 73% of gene editing efficiency. The other edited C led to a silent mutation GAC (D) to GAT (D). (D) Q94* mutation in Tek upon C-to-T conversion in *tek* reached 18% of gene editing efficiency. (E) Q273* mutation in Bap1 upon C-to-T conversion in *bap1* reached 14% of gene editing efficiency. (F) Q21* mutation in p53 upon C-to-T conversion in *tp53* reached 73% of gene editing efficiency. (G) Q170* mutation in p53 upon C-to-T conversion in *tp53* reached 86% of gene editing efficiency. For (C) and (E) The reverse complement of the sgRNA sequence is shown. Scale bars: (B) 50μm.

Here based on these technological advances, we optimized these second-generation gene editing tools for the zebrafish in order to overcome limitations of HDR-based approaches in this organism. As reported in *ex vivo* studies (Koblan et al., 2018), we tested different CBE variants and obtained highly efficient C-to-T conversion, reaching up to 91% efficiency without unwanted mutations. Furthermore, we used these tools to target Wnt signaling, thus proving that endogenous pathways can be modulated in their natural context. Finally, we demonstrate the power of this technology for introducing precise mutations in human pathology-associated genes with high efficiency in zebrafish.

## RESULTS AND DISCUSSION

### BE4-gam base editing for the endogenous activation of Wnt signaling pathway

To date, the main strategies used in zebrafish to study the constitutive activation of signaling pathways and to dissect their role during embryonic development or tumorigenesis have been based on overexpressing mutated genes. To gain more insights and to complement these studies, an important requirement is to have the ability to maintain the endogenous genetic and regulatory contexts by generating mutations of endogenous genes *in vivo*.

To address this challenge, we decided to introduce an activating mutation in the *ctnnb1* gene coding for the key effector β-catenin of canonical Wnt signaling, a major signaling pathway during embryonic development which is activated in many cancers (Steinhart and Angers, 2018). It was previously shown that mutation of Serine33 into a Leucine of the human β-catenin protein prevents its degradation by the ubiquitin-proteasome system, leading to its stabilization, thereby constitutively activating Wnt signaling (Hart et al, 1999) (Liu et al., 1999).

We first aimed at introducing this mutation in the genome of the zebrafish by using the BE4-gam CBE (Fig. 1A). This CBE was indeed one of the first variants of CBEs to show high efficiency of gene editing and fewer INDELs formation in cultured cells (Komor et al., 2017). We injected the *BE4-gam* mRNA and synthetic *ctnnb1 S33L* sgRNA into one-cell stage *Tg(tcf:gfp)* zebrafish embryos to directly monitor the effect of the introduced mutation on the activity of the canonical Wnt signaling (Moro et al., 2012). Upon *ctnnb1 S33L* sgRNA injection, we observed an increase of GFP-positive cells at 1dpf compared to the control embryos, as well as an ectopic activation of the pathway in retinal progenitor cells (Fig. 1B). Using this strategy, we were able to reach up to 73% of editing efficiency. In addition, we also observed conversion of another cytidine within the PAM [−19, −13 bp] window leading to a silent mutation (GAC-to-GAT (D)) with up to 74% efficiency (Fig. 1C, Table.1, supplementary Table.3).

**Table.1.**
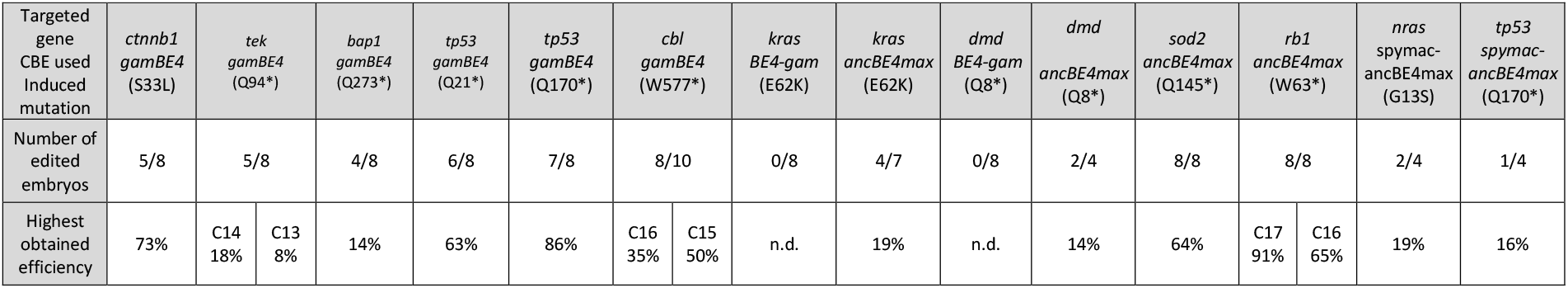
Base-editing efficiency using different CBE variants. Number of edited embryos randomly chosen after injection of *CBE* mRNA and sgRNA. The efficiency varies between non-detected (n.d.) and 91% depending on the targeted locus, the sgRNA and the CBE used. Editing efficiency was quantified by editR analysis(Kluesner et al., 2018) which does not detect editing efficiency below 5%.

With these results we demonstrated that it is now possible to constitutively and efficiently activate important developmental signaling pathways in their endogenous context, as we show here for Wnt signaling. Furthermore, several studies have implicated the S33L β-catenin mutation in tumorigenesis, making it possible to study the role of this oncogenic mutation in cancer development in zebrafish. In order to test the potential of CBE targeting in cancer modelling, we then decided to use it to target a series of tumor suppressor genes and oncogenes using the same editing strategy applied to endogenous β-catenin.

### Base editing strategies for the generation of human cancer mutations

Zebrafish is a powerful model system to study cancer genetics *in vivo* (Cagan et al., 2019; Cayuela et al., 2018). However, a robust method for modeling cancer-associated mutations in zebrafish is lacking to date. We decided to create predictable premature stop codons in tumor suppressor genes and to generate activating-mutations in oncogenes of the RAS family (Li et al., 2018) in order to test the ability of CBE to induce cancer-related mutations in zebrafish. We first developed an automated script to rapidly detect codons allowing to generate nonsense mutations after C-to-T conversion within the restricted PAM [−19, −13] bp window of edition (sequenceParser.py). Using this script, we designed a series of sgRNAs targeting in a selection of tumor suppressor genes. We could induce the Q94* mutation in Tek, Q273* mutation in Bap1 and Q21* mutation in p53 as well as Q170* in p53 by C-to-T conversions (Fig. 1D-G, Table.1, supplementary Table.3). Among the different targeted mutations, the highest efficiency achieved was for the *tp53* tumor suppressor gene, for which we reached up to 86% of C-to-T conversion for the introduction of the Q170* mutation (Fig. 1G, Table.1, supplementary Table.3). This last result is particularly remarkable as the best conversion rate obtained using base editing in zebrafish to date was a maximum of 28% efficiency with an INDEL score of 5% using the BE3 variant (Zhang et al., 2017). To assess the presence of INDELs or unwanted mutations upon BE4-gam injections in our targets, we amplified, cloned and sequenced all targeted loci. For *tek* 6 out of 20, *bap1* 2 out of 12, *tp53* (Q21*) 12 out of 24 and lastly *tp53* (Q170*) 21 out of 24 colonies showed precise C-to-T conversion whereas all the other analyzed sequences were wild-type, without any error or INDEL formation.

Together these results show that, using BE4-gam, we could efficiently target several genes implicated in tumorigenesis in zebrafish without generating any unwanted INDELs, unlike what had been previously reported with the BE3 variant.

More recently a new CBE variant, the ancBE4max, has been engineered and optimized in cell culture with increased efficiency compared to the classical BE4-gam, reaching up to 90% efficiency and very low rates of INDELs (Koblan et al., 2018). We therefore decided to use this new CBE variant to target the oncogenic mutation E62K in Kras and induce the creation of a Q8* stop codon in the Dmd tumor suppressor for which we were unsuccessful using the BE4-gam (Table1, supplementary Table.3). By co-injection of *ancBE4max* mRNA with the *kras E62K* sgRNA, we were able to introduce the E62K mutation with up to 19% of efficiency. Another cytidine in the editing window was also converted and led to the generation of a silent mutation (CAG-to-CAA (Q)) (Fig. 2A, Table.1, supplementary Table.3). Similar to what we observed in the case of *kras* editing, we were able to obtain a Q8* mutation in the Dmd tumor suppressor (Fig. 2B, Table.1, supplementary Table.3). Thus, with this new ancBE4max variant, we are able to introduce mutations that could not be achieved with BE4-gam using the same sgRNAs. Remarkable editing efficiency was also observed using this CBE for two additional targets: the tumor suppressor genes *sod2* and *rb1*, for which respectively up to 64% and 91% of editing were reached and 100% of the sequenced embryos were precisely mutated (Fig. 2C, D, Table.1, supplementary Table.3) (Bravard et al., 1992; Dyson, 2016).

**Fig. 2.**
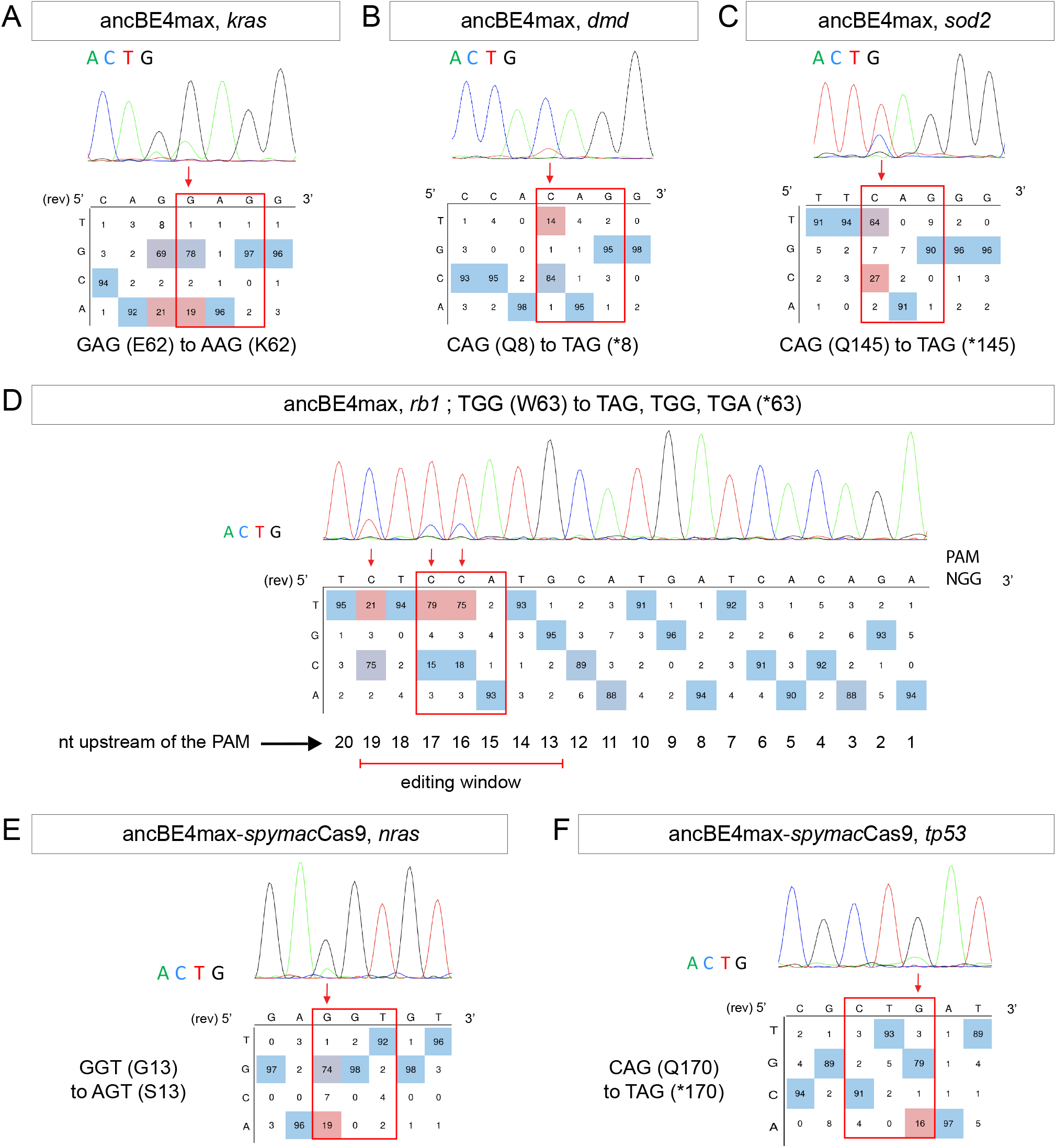
Tumor suppressor genes and oncogenes targeting by the highly efficient ancBE4max and the *ancBE4max-SpymacCas9* recognizing NAA PAM. (A-F) DNA sequencing chromatogram of targeted loci with the ancBE4max (in A-D) or ancBE4max-*Spymac*Cas9 (in E-F) and obtained C-to-T conversion efficiencies. (A) E62K mutation in Kras upon C-to-T conversion in *kras* reached 19% gene editing efficiency. The other edited C led to a silent mutation CAG (Q) to CAA (Q). (B) Q8* mutation in Dmd upon C-to-T conversion in *dmd* reached 14% of gene editing efficiency. (C) Q145* mutation in Sod2 upon C-to-T conversion in *sod2* reached 64% of gene editing efficiency. (D) W63* mutation in Rb1 upon C-to-T conversion in *rb1* reached 21% for the C19 base, 79% for C17 and 75% for the C16 of gene editing efficiency. (E) G13S mutation in Nras upon C-to-T conversion in *nras* reached 19% of gene editing efficiency. (F) Q170* mutation in p53 upon C-to-T conversion in *tp53* reached 16% of gene editing efficiency. For (A, D-F) The reverse complement of the sgRNA sequence is shown. (A-F) It is considered that the quantifications under 5% are due to the background signal from Sanger sequencing and are thus non-significant(Kluesner et al., 2018).

It is interesting to note that in general all cytidine bases in the PAM [−19, −13] bp window can be edited by the CBE, with a higher efficiency for the cytidine bases located in the middle of this window while editing was below detection levels for cytidines located only 12bp upstream (Fig. 2D). With the use of ancBE4max CBE, these results highlight the importance of the cytidine distance from the PAM for efficient editing in zebrafish as shown previously in cell culture assays (Gaudelli et al., 2017).

### Expanding new gene editing possibilities in zebrafish using a CBE recognizing NAA PAM

Due to the PAM-dependent restriction of the editing window, many mutations could not be achieved so far. We therefore decided to expand the editing possibilities in zebrafish by associating *SpymacCas9* recognizing NAA PAMs with the efficient conversion capacity of the ancBE4max. To this end, we replaced the PIM domain (PAM-Interacting Motif) of the *Sp*Cas9 with the one of the *SpymacCas9* in the ancBE4max (Jakimo et al., 2018). The inserted PIM domain was codon optimized for zebrafish. Using this newly generated ancBE4max-*SpymacCas9*, we were able to reproduce the human G13S mutation in *nras* oncogene in zebrafish with up to 19% of efficiency (Fig. 2E, Table.1, supplementary Table.3). We also introduced a stop codon by a C-to-T conversion in the *tp53* gene with up to 16% efficiency (Fig. 2F, Table.1, supplementary Table.3). These results demonstrate that in addition to the classical NGG PAM it is now also possible to target NAA PAMs in zebrafish, thereby significantly expanding the range of cytidine bases that can be converted. For these new CBEs, we added a function in our script to choose the PAM recognized by the Cas9 of the chosen CBE to generate the desired editing (sequenceParser.py). Together with the use of the ancBE4max and *ancBE4max-SpymacCas9* CBE variants, we were now able to target mutations that could not be generated with the BE4-gam base editor and reproduce a wider range of human cancer mutations in zebrafish.

Genetic alterations that lead to oncogene activation and/or tumor suppressor inactivation are responsible for tumorigenesis. It is indeed well established that in cancer patients, a series of genetic mutations in tumor suppressor genes and/or oncogenes are combined to all together lead to the appearance of the disease (Dash et al., 2019). With these efficient genetic tools that are now established in zebrafish, we have the possibility to rapidly test precise combinations of mutations identified in cancer patients.

### Precise gene editing in *cbl* for the generation of human disease phenotypes in zebrafish

With the technological advances on CRISPR/Cas9 gene editing, zebrafish has become an even more attractive system for modeling human genetic diseases. Among the loci chosen to test the efficiency of the BE4-gam, we targeted the tumor suppressor gene encoding for Cbl, a E3 ubiquitin ligase, that is found mutated in Noonan syndrome patients presenting short stature and other bone malformations among other phenotypes (Martinelli et al., 2010). In humans, activating mutations in the *fibroblast growth factor receptor 3 (FGFR3)* gene are a leading cause of dwarfism achondroplasia and related dwarfing conditions as FGFR3 hyperactivation triggers intracellular signaling within the chondrocytes of the growth plate that terminates its proliferation and bone growth (Harada et al., 2009). Interestingly, another study based on *in vitro* systems reported that some of these activating mutations in *FGFR3* disrupt c-Cbl-mediated ubiquitination that serves as a targeting signal for lysosomal degradation and termination of receptor signaling (Cho et al., 2004). Using the CBE BE4-gam as previously described, we obtained up to 50% of gene editing efficiency for zebrafish *cbl* (Fig. 3A, Table.1, supplementary Table.3) with 80% of the analyzed embryos showing the expected editing. 4 out of 15 adults carried the Cbl W577* mutation in germ cells and transmitted it to 28% of their F1 offspring (44 out of 153 analyzed fish carried the mutation). The target sequence was analyzed in the F1 embryos and no INDELs were found (Fig. 3B). Interestingly, 24% of the maternal zygotic *MZ cbl*^-/-^ mutants displayed a significant reduced overall growth and size by 3 months post-fertilization while 100% of the wild-type sibling fish showed a normal body size (means: 2,7cm for the wild-type siblings *MZ cbl*^+/+^ and 1,96cm for *MZ cbl^-/-^*, Fig. 3C, D). Four germline mutations located in the RING domain of Cbl (Q367P, K382E, D390Y and R420Q) have been previously identified and associated to Noonan Syndrome and related phenotypes (Martinelli et al., 2010). Our results are in line with the growth defect phenotypes observed in these patients and directly implicate Cbl loss-of-function as a cause of bone malformations in an animal model. In addition, a point mutation in zebrafish Cbl (H382Y) has been implicated in myeloproliferative disorders. Unlike our mutant, *cbl^H382Y^* mutants do not survive to adulthood, suggesting that the Cbl^W577*^ premature stop reported here may have different consequences on the multiple functions of Cbl (Peng et al., 2015). Although not lethal, it would be of interest to assess whether any hematopoietic defects are present in our *MZ cbl*^-/-^ mutants or whether this phenotype is only linked to the Cbl^H328Y^ substitution found in the *LDD731* zebrafish mutant (Peng et al., 2015). Our model represents a powerful *in vivo* system to dissect the role of *cbl* in bone morphogenesis and to explain the human phenotype related to bone malformations.

**Fig. 3.**
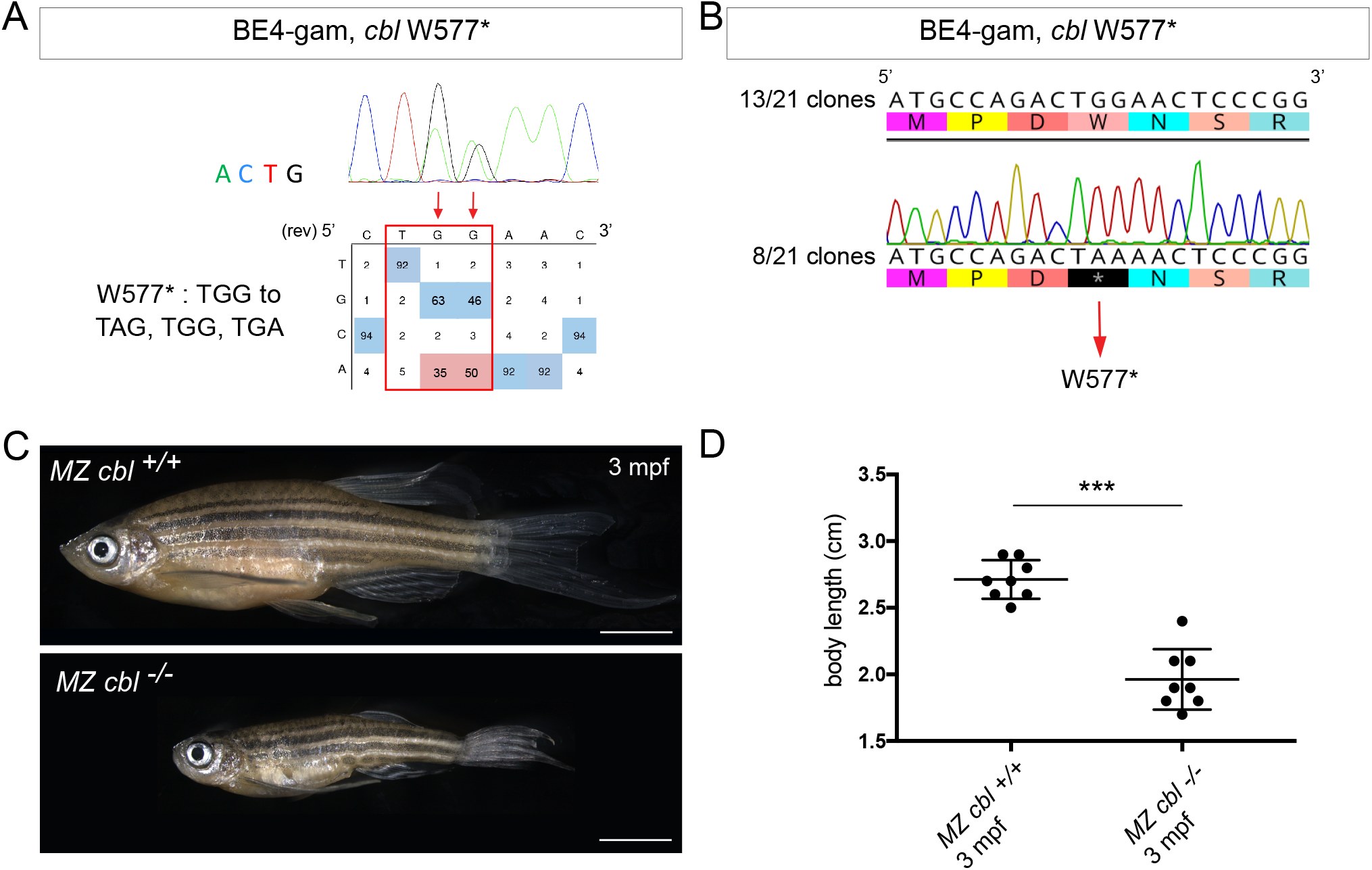
BE4-gam generated *cbl* maternal zygotic mutant fish show a reduced growth phenotype. (A) DNA sequencing chromatogram of targeted *cbl* gene with the BE4-gam. W577* mutation in Cbl upon C-to-T conversion in *cbl* reached 50% for the C16 base and 35% for the C15 base of gene editing efficiency. It is considered that the quantifications under 5% are due to the background signal from Sanger sequencing and are thus non-significant(Kluesner et al., 2018). (B) Sequencing of individual clones of a pool of F1 embryos from a founder carrying the W577* mutation in Cbl. TGG-to-TAA precise mutation was found in 8 out of 21 clones. No edition nor INDELs were detected in all other clones. (C) 3 months post-fertilization (mpf) *cbl* maternal zygotic wild type sibling (upper panel) and mutant fish (lower panel). (D) 24% of *MZ cbl^-/-^* mutant fish shows a significant reduced size at 3 mpf compared to wild-type siblings as shown by the quantification of their respectively body length. n=8 for each group. Mann-Whitney test, p<0,0005. Scale bars: (C) 5mm.

## Conclusions

In our work, we took advantage of base editors to generate C-to-T point mutations at unprecedented high efficiencies (up to 91%) without detecting any unwanted mutations that were so far problematic when using CBEs in zebrafish. To expand the gene editing possibilities in this animal model, we established a new editor variant recognizing the NAA PAM. Using these approaches, we first performed the endogenous and constitutive activation of Wnt signaling by introducing the S33L mutation in β-catenin. In addition, we demonstrated using these strategies that we were able to precisely target several cancer-associated genes for which so far only transgenic over-expressions or imprecise deletions were used to elucidate their functions. Among our targets, the introduced mutation in the *cbl* gene, coding for a E3 ubiquitin ligase, allowed us to generate a new zebrafish model for dwarfism.

Together our work provides a panel of examples whereby, using gene editing approaches, some of which established here in zebrafish for the first time, it is now possible to manipulate endogenous signaling pathways, generate models for human genetic disorders and mimic precise cancer-associated mutations in zebrafish. While this manuscript was in preparation, a study was reported where the authors used the CBE system in zebrafish modelled the pathological features of human ablepharon macrostomia syndrome (AMS) (Zhao et al., 2020). In line with our findings, such study highlights the power and the need for these approaches in order to model pathological human mutations in zebrafish.

Finally, the high efficiencies of CBEs obtained in this study will permit future applications where they could be implemented with mosaic mutation induction technologies such as the MAZERATI (Modeling Approach in Zebrafish for Rapid Tumor Initiation) system (Ablain et al., 2018). In the future, this will allow to rapidly model and study *in vivo* combinations of endogenous mutations occurring in specific cancer patients or in genetic disorders caused by somatic mosaicism. Our approach could thus be applied in zebrafish for the precise modelling of complex combinations of cancer-causing mutations in adult animal models as currently possible by transgenic overexpression or somatic gene inactivation (Callahan et al., 2018).

## Acknowledgements

We thank Sophie Vriz for sharing the *Tg(tcf:gfp)* transgenic line and the members of the fish-facility in Institut Curie. We also thank Céline Revenu and Viviana Anelli for early contribution. M.R. was supported by the Fondation pour la Recherche Médicale (FRM grant number ECO20170637481) and la Ligue Nationale Contre le Cancer. Work in the Del Bene laboratory was supported by ANR-18-CE16 “iReelAx”, UNADEV in partnership with ITMO NNP/AVIESAN (national alliance for life sciences and health) in the framework of research on vision and IHU FOReSIGHT [ANR-18-IAHU-0001] supported by French state funds managed by the Agence Nationale de la Recherche within the Investissements d’Avenir program. M.C.M. was supported by World Wide Cancer Research, grant no. 0624, and by LILT –Trento, Program 5 per mille (year 2014).

## Methods

### Fish lines and husbandry

Zebrafish (Danio rerio) were maintained at 28 °C on a 14 h light/10 h dark cycle. Fish were housed in the animal facility of our laboratory which was built according to the respective local animal welfare standards. All animal procedures were performed in accordance with French and European Union animal welfare guidelines. Animal handling and experimental procedures were approved by the Committee on ethics of animal experimentation. *Tg(tcf:gfp)* was kindly provided by Sophie Vriz(Moro et al., 2012).

### Molecular cloning

To generate the *pCS2+_ancBE4max-SpymacCas9* plasmid, the *SpymacCas9* PIM domain sequence has been codon optimized for expression in zebrafish using online software from IDT and synthesized with the first UGI sequence as G-block from IDT. Then, 3 fragments have been inserted into *pCS2+ plasmid* linearized with Xho1 using the Gibson Assembly Cloning Kit (New England Biolabs): a first fragment of 4161bp of the ancBE4max to the PIM domain (amplified using the primers F-5’-CGATTCGAATTCAAGGCCTCATGAAACGGACAGCCGAC-3’ and R-5’-CGGTCTGGATCTCGGTCTTTTTCACGATATTC-3’), the Gblock fragment of 803bp (amplified using the primers F-5’-AAAGACCGAGATCCAGACCGTGGGACAG-3’ and R-5’-TCCCGCCGCTATCCTCGCCGATCTTGGAC-3’) and a third fragment of 654bp of the rest of the ancBE4max from the PIM domain (amplified using the primers F-5’-CGGCGAGGATAGCGGCGGGAGCGGCGGG-3’ and R-5’-CTCACTATAGTTCTAGAGGCTTAGACTTTCCTCTTCTTCTTGGGCTCGAATTCGCTGCCGTCG-3’).

### mRNAs synthesis

*pCMV-BE4-gam* (a gift from David Liu, Addgene plasmid # 100806)(Anzalone et al., 2019) have been used to generate *BE4-gam in vitro*. These plasmids were linearized with Pme1 restriction enzyme and mRNAs were synthesized by *in vitro* transcription with 1μl of GTP from the kit added to the mix, followed by Poly(A) tailing procedure and lithium chloride precipitation (using the mMESSAGE mMACHINE T7 Ultra kit #AM1345, Ambion). *pCMV-ancBE4max* (pCMV_AncBE4max was a gift from David Liu (Addgene plasmid # 112094) has been linearized using AvrII restriction enzyme, mRNAs were synthesized by *in vitro* transcription with 1μl of GTP from the kit added to the mix and lithium chloride precipitation (using the mMESSAGE mMACHINE T7 Ultra kit #AM1345, Ambion). the *pCS2+_ancBE4max-SpymacCas9* has been linearized using KpnI restriction enzyme, mRNAs were synthetized by *in vitro* transcription with 1μl of GTP added to the mix and lithium chloride precipitation (using the mMESSAGE mMACHINE sp6 Ultra kit #AM1340, Ambion).

### sgRNA design

A sequenceParser.py python script was developed and used to design sgRNAs for the creation of a stop codon. The first function of the script is to ask which PAM will be used to then execute the rapid detection of codons that are in the right editing windows from this predefined PAM to generate a STOP in frame after C-to-T conversion. The ORF sequence file extension is .txt and the letters in lower cases. The script can be executed from the command line interface (CLI, such as the terminal or PowerShell console).

Efficiencies of sgRNAs were validated using CRISPOR online tool (Haeussler et al., 2016). All the synthetic sgRNAs were synthesized by IDT.

### Micro-injection

To make the mutagenesis with base editing, a mix of *ancBE4max* mRNA or *BE4-gam* mRNA and a synthetic sgRNA was injected in one-cell stage zebrafish embryos. The final concentration was 600ng/μL for *CBE* mRNA and 43pmol/μL for sgRNA.

### Genotyping

To genotype the *cbl* mutant line, a PCR was performed with primers Fwd-5’-GTACGCCTGGAGACCCATCTC-3’ and Rev-5’CTTTTGGACTGTCATAATCCGATGC-3’. The PCR product was digested with the restriction enzyme BsrI, which cut only on the WT allele. The WT allele resulted in two fragments (300bp and 69bp) and the mutant allele only one fragment (369bp).

### Whole-embryos DNA sequencing

A series between 4 and 10 single embryos randomly chosen was analyzed for each target sequence and the embryo with the highest efficiency is shown. Generally, between 25% to 100% embryos were positive for gene editing, i.e. showed >16% expected sequence modification. For genomic DNA extraction, each single embryo was digested for 1 h at 55°C in 0.5 mL lysis buffer (10 mM Tris, pH 8.0, 10 mM NaCl, 10 mM EDTA, and 2% SDS) with proteinase K (0.17 mg/mL, Roche Diagnostics) and inactivated 10 min at 95°C. To sequence and check for frequency of mutations, each target genomic locus was PCR-amplified using Phusion High-Fidelity DNA polymerase (Thermo Scientific). For the base editor mutations, the PCR product was sent for sequencing, analyzed using ApE software and quantifications of the mutation rate done using editR online software(Kluesner et al., 2018). For the verification of *cbl* mutant F1 embryos and *tp53* mutation, PCR fragments were subsequently cloned into the pCR-bluntII-TOPO vector (Invitrogen). Plasmid DNA was isolated from single colonies and sent for sequencing. Mutant alleles were identified by comparison with the wild-type sequence using ApE and Geneious softwares.

### Imaging

Embryos were oriented in low melting agarose 0,6% with an anesthetic (Tricaine 0,013%) diluted in egg solution. The inverted laser scanning confocal microscope Zeiss CLSM - LSM780 was used for high resolution microscopy, employing a 40x water immersion objective. Z-stacks were acquired every 1- to 2-mm. Leica MZ10F was used to image the whole embryos the *cbl* mutant adult fish. Image analyses were performed with ImageJ software.

### Body size quantifications

8 *MZcbl*^+/+^ wild-type siblings and 8 *MZcbl*^-/-^ in total were used to measure the body size using a millimetric ruler. The length measured was from mouth to trunk. A non-parametric t-test with the Mann-Whitney correction was applied to determine significance in growth. The software used was Prism 7 (GraphPad).

---

**Supplementary Table.1.**
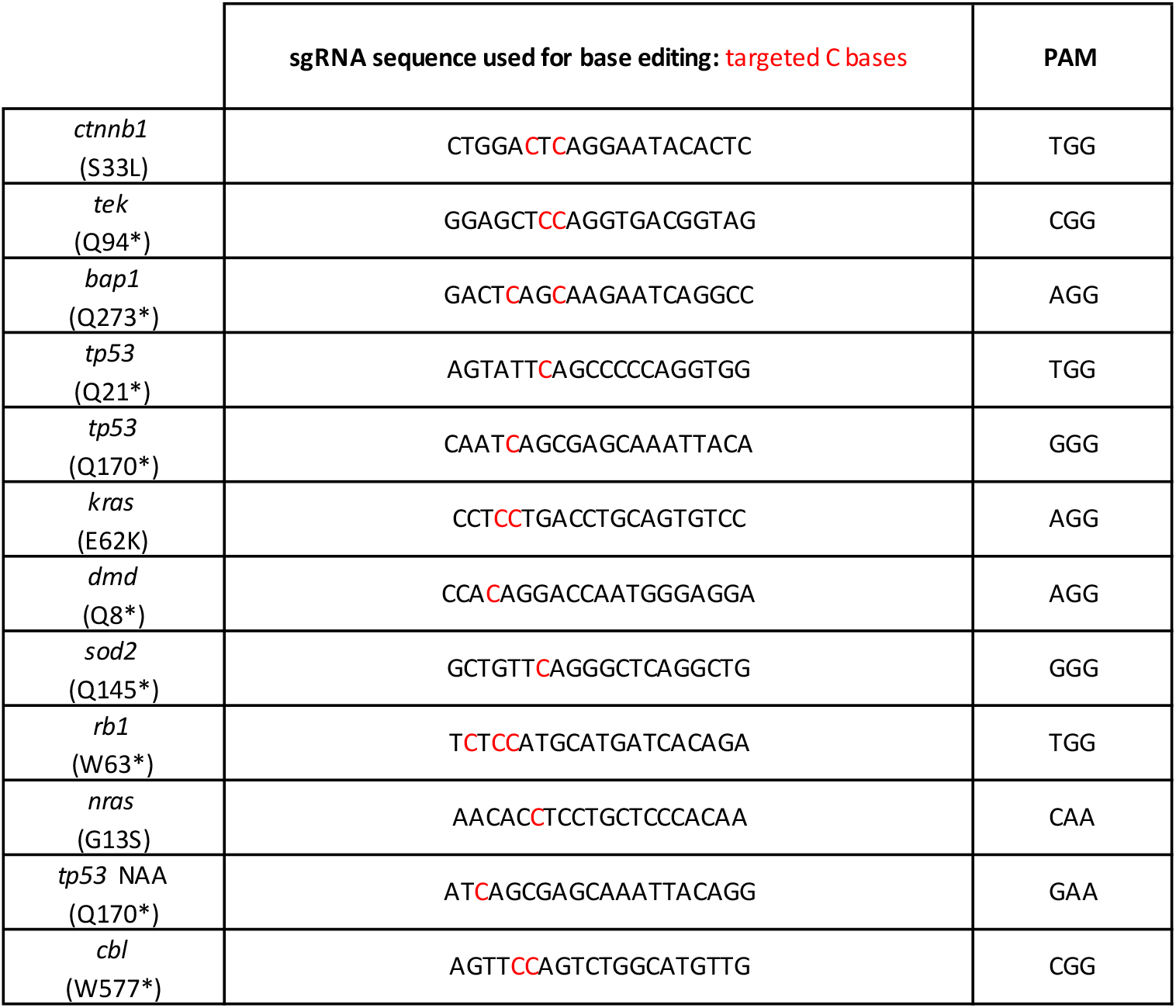
sgRNA sequences.

**Supplementary Table.2.**
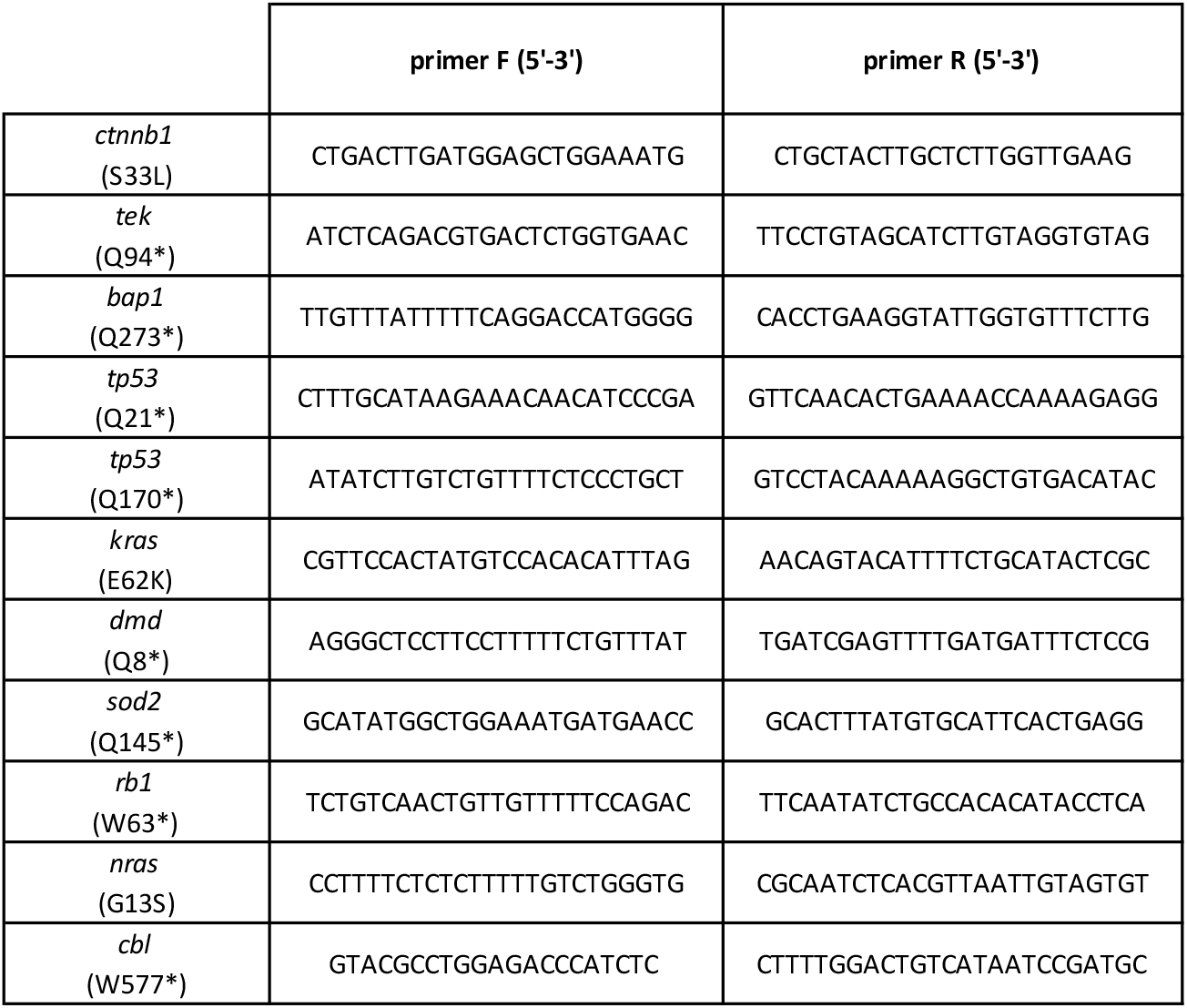
Primers sequences used to amplify the targeted loci.

**Supplementary Table.3.**
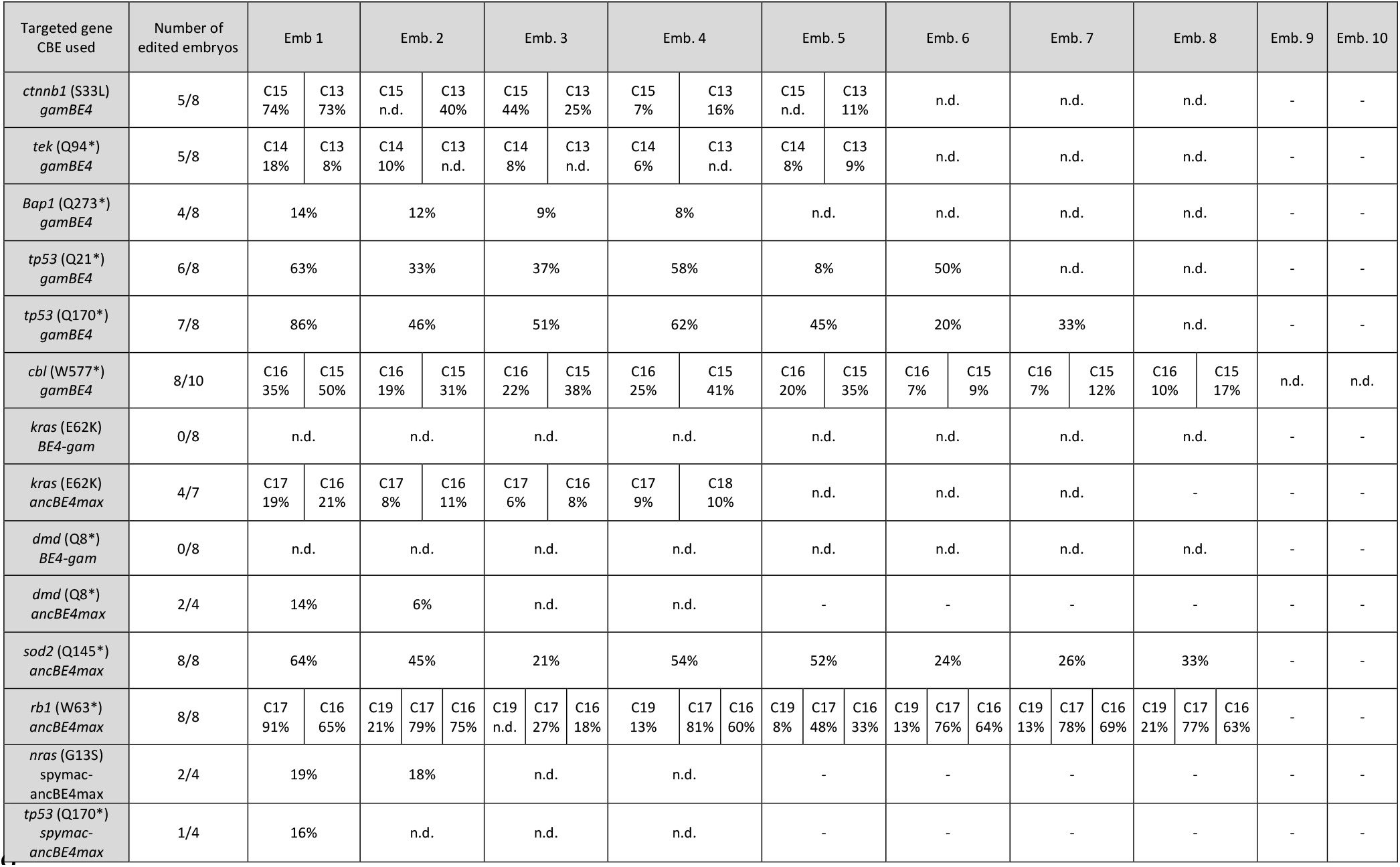
Editing efficiency quantification. Editing quantification of up to 10 single embryos randomly chosen after injection of *CBE* mRNA and sgRNA. The efficiency varies between non-detected (n.d.) and 91% depending on the targeted locus, the sgRNA and the CBE used. Editing efficiency was quantified by editR analysis(Kluesner et al., 2018) which does not detect editing efficiency below 5%.

